# A comparative study of cellular diversity between the *Xenopus* pronephric and mouse metanephric nephron

**DOI:** 10.1101/2022.01.11.475739

**Authors:** Mark E. Corkins, MaryAnne Achieng, Bridget D. DeLay, Vanja Krneta-Stankic, Margo P. Cain, Brandy L. Walker, Jichao Chen, Nils O. Lindström, Rachel K. Miller

## Abstract

The kidney is an essential organ that ensures bodily fluid homeostasis and removes soluble waste products from the organism. The functional units within the kidneys are epithelial tubules called nephrons. These tubules take in filtrate from the blood or coelom and selectively reabsorb nutrients through evolutionarily conserved nephron segments, leaving waste product to be eliminated in the urine. Genes coding for functional transporters are segmentally expressed, enabling nephrons to function as selective filters. The developmental patterning program that generates these segments is of great interest. The *Xenopus* embryonic kidney, the pronephros, has served as a valuable model to identify genes involved in nephron formation and patterning. Prior work has defined the gene expression profiles of *Xenopus* epithelial nephron segments via *in situ* hybridization strategies, but our understanding of the cellular makeup of the *Xenopus* pronephric kidney remains incomplete. Here, we scrutinize the cellular composition of the *Xenopus* pronephric nephron through comparative analyses with previous *Xenopus* studies and single-cell mRNA sequencing of the adult mouse kidney, this study reconstructs the cellular makeup of the pronephric kidney and identifies conserved cells, segments, and expression profiles. The data highlight significant conservation in podocytes, proximal and distal tubule cells and divergence in cellular composition underlying the evolution of the corticomedullary axis, while emphasizing the *Xenopus* pronephros as a model for physiology and disease.

**Summary Statement:** Assessment of the conservation of gene expression between functional *Xenopus laevis* pronephric and mouse metanephric nephrons through single-cell RNA sequencing.

## Introduction

The kidney is an essential organ that maintains organismal fluid homeostasis and waste excretion. It develops in three successive forms, the pronephros, mesonephros and metanephros, with each new stage displaying increased architectural complexity ^1^. The pronephros, the embryonic kidney, consists of a single epithelial tubule called a nephron, providing the structural and signaling foundation for the mesonephros to build upon. Additional nephrons form along the nephric duct that elongates from the pronephros, establishing the mesonephros. The nephric duct or Wolffian duct is also the foundation of the male reproductive system. The mesonephros is the adult kidney stage in amphibians and fish. In amniotes, such as birds, reptiles, and mammals, a third kidney called the metanephros forms upon branching of the ureteric bud from the nephric duct, replacing the function of the mesonephros. All three kidney stages utilize nephrons to filter blood or coelomic fluid. Nephrons reabsorb nutrients that the organism needs, leaving wastes to be excreted in the urine. The nephron is structured to reabsorb substances within specific regions along the nephron, and genes expressed in each segment enable distinct functions ^2–5^. The pro, meso, and metanephric kidneys reuse genetic pathways to generate new nephrons, conserving these structures through each successive stage of kidney development. Disruption of genetic pathways involved in the specification or patterning of the kidney lead to a number of kidney diseases ranging from kidney hypoplasia, where parts of the kidney do not develop, to kidney agenesis, where the kidney fails to form ^6–9^. Additionally, other kidney diseases preferentially manifest within specific nephron segments, such as polycystic kidney disease ^10–12^. Understanding the cell types and developmental programs underpinning nephron patterning will provide insights into global mechanisms underlying kidney disease.

*Xenopus* is an informative model in which to study kidney development ^13^. The *Xenopus* pronephric kidney consists of a single nephron ^14^, which is visible through the organism’s semitransparent epidermis. This kidney’s simplicity enables visualization of processes essential to kidney development and *in vivo* imaging ^15–17^. Additionally, the ability to quickly knock out, knock down, or overexpress genes in *Xenopus* has facilitated a mechanistic understanding of kidney development. Though many factors regulating nephrogenesis are conserved between *Xenopus* and humans ^6,18^, the extent of this conservation has not been extensively evaluated.

Previous studies have evaluated the cellular makeup and spatial patterning of gene expression within *Xenopus* kidneys by evaluating physiological structures and mRNA expression patterns ^14,19–22^. These studies provided valuable insight into the spatial organization of the kidney. However, relatively few cells are physiologically distinguishable from each other, and the number of genes evaluated was limited by technical challenges of scaling *in situ* hybridization analysis. The use of colorimetric immunohistochemistry staining for the detection of *in situ* probes does not allow cellular resolution of gene expression. Because studies in other model organisms relied on similar techniques, the cellular and transcriptional makeup for a reference kidney has also been incomplete. This has resulted in the development of multiple distinct models of the patterning of the *Xenopus* kidney.

In recent years, mRNA-Seq advances have facilitated the sequencing of individual cells ^23^. The *Xenopus* model organism is advantageous for single-cell analysis given that the embryo undergoes holoblastic cleavage, and each cell contains sufficient nutrients in the form of yolk to allow cells to be easily cultured. In addition, dissociation and culture requirements have been optimized for several embryonic cell types ^15,24–26^, making *Xenopus* a suitable organism for primary culture and single-cell sequencing experiments. In *Xenopus tropicalis* single-cell mRNA sequencing experiments have been carried out ^27–29^. In *Xenopus laevis*, the only single-cell experiment published thus far evaluated tail regeneration ^30^, possibly due to the challenges associated with ^31^evaluating transcripts derived from its allotetraploid genome. Recently, multiple single-cell sequencing experiments have been carried out in mouse ^32–34^ and human kidneys ^31,35,36^. Therefore, the mouse kidney can now be used as a reference to identify conserved cell types in the *Xenopus* kidney.

To identify the cellular composition of the *Xenopus* kidney, we performed single-cell mRNA sequencing on kidneys dissected from the allotetraploid *X. laevis*. Using previous studies showing the spatial expression patterns in the *Xenopus* nephron along with single-cell mRNA sequencing data from the adult mouse kidney, we analyzed and reconstructed the cellular makeup of the pronephric kidney.

## Results

### Cellular diversity in the Xenopus pronephros

The *Xenopus* embryonic kidney starts forming during neurulation (stage 12.5), and the nephron is fully functional by tadpole stage (stage 40) ^22^. To scrutinize the cellular composition of the functional *Xenopus* nephron, pronephroi were manually dissected from approximately 300 stage 40 *X. laevis* embryos (Movie S1) and processed for single-cell mRNA sequencing. The viability of the dissociated cells was assessed by trypan blue staining, and the mRNA from living cells was sequenced using the 10X Genomics platform (v3 Chemistry). Sequencing results were obtained from 6,476 cells with an average of 23,515 reads per cell. The sequence reads were compared with the *X. laevis* prerelease genome (Jan 2020) (Table S1). The sex of the cells sequenced from *Xenopus* nephrons were indistinguishable based on commonly used markers (Dmrt1, Foxl2, Dm-w), which were not detected in our samples ^37^ Therefore sex was ignored.

Because *X. laevis* has an allotetraploid genome resulting from a whole genome fusion approximately 17-18 million years ago, most genes have two homeologs, labeled L (Long Chromosome) and S (Short Chromosome)^38^, which were analyzed separately (Fig. S1, S2). Given that the genes produced from both homeologs similar in sequence, techniques such as *in situs* are not often designed distinguish between homeologs. Therefore, to facilitate the comparison of these homeologs to the diploid mouse transcripts, these transcripts were merged (Fig. 1–6, S3). The L and S versions of the gene share the same name; therefore, we combined homeolog pairs into a single gene by summing raw gene counts for each (Table S2, S3). The combined counts were used for this analysis unless otherwise stated.

**Figure 1:**
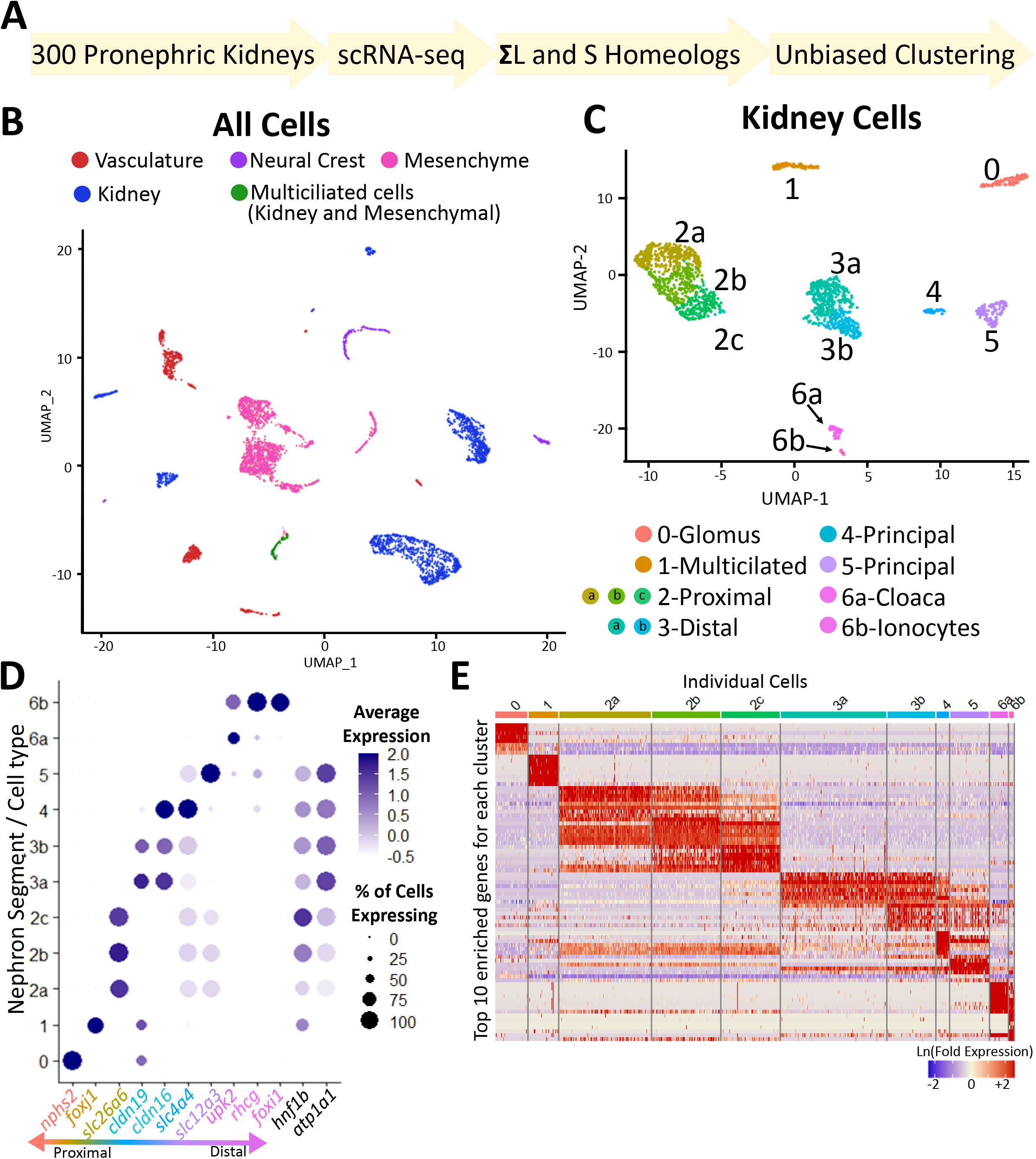
Single-cell analysis of dissected kidneys from stage 40 *X. laevis*. **A.)** Description of methodology to generate integrated data sets. **B.)** UMAP showing the cellular makeup of all the cells isolated along with the *X. laevis* kidney. Multiciliated cells are represented by green cells as they are present in both the kidney and mesenchymal cell types. **C.)** UMAP showing the kidney subset of cells. Clusters were numbered from most proximal to most distal. **D.)** Dotplot showing the expression of genes previously identified by *in situ* analysis for each region of the kidney. **E.)** Heatmap showing the expression of the top 10 most enriched genes for each cluster. Rows indicate gene expression and columns indicate a single cell.

### Single-cell profiling and clustering of Xenopus kidney cells

Kidney cells were identified by the expression of known kidney specific markers (*cdh1, cdh16, nphs2, atp1a1, hnf1b*) and analyzed independently of other isolated cell types (Fig. 1B and S1B). UMAP projection of the kidney single-cell sequencing data clearly separates the cells into seven independent clusters/cell types (Fig. 1C and S1C). Using markers defined by Raciti et al., 2008; Reggiani et al., 2007; Zhou and Vize, 2004, we numbered and color-coded the segments in order from proximal to distal along the nephron (Fig. 1C,D and S1C,D). Cluster segments were validated by visualizing gene expression for the top ten enriched genes within each of the identified cell types (Fig. 1E, S1E).

Cells such as those within the proximal and distal early nephron segments separated by unbiased clustering, but they remain grouped as closely related types on the UMAP projection (Fig. 1C and S1C). Closely related clusters were labeled with a letter such as 2a-c. Though these clusters had unique genes, few transcripts were completely restricted to one subtype (Fig. 1E, S1E, and S3). By comparing the enriched genes from each group of the proximal tubule (2a-c) and their known localization, as determined by *in situ* analyses ^5,21,22^, we found that each cluster likely correlated to the three different sections of the proximal tubules (Fig 2A). Similar analyses were performed for distal cells (3-5), which also corresponded to known distal cell types within anatomically distinct regions (Fig 2B).

**Figure 2:**
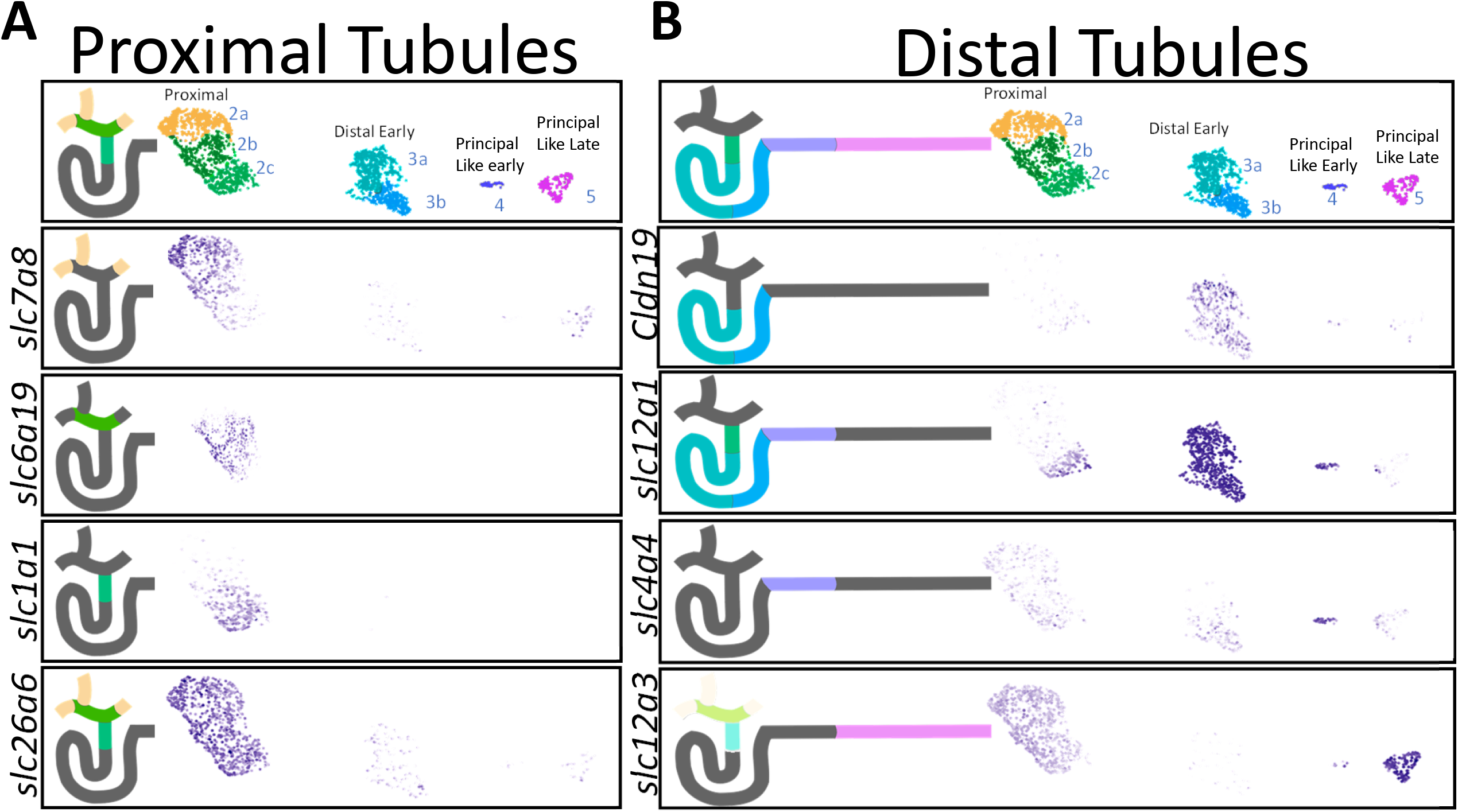
Comparison of nephron cell types with *X. laevis in situ* patterning. Clusters indicate special localization of gene expression in the proximal **(A)** and distal **(B)** tubules. Diagram shows expression of gene as identified by *in situ* analysis from ^21^). Colored regions indicate expression.

Cluster 6 initially remained as a single cluster, though the UMAP projection generated two independent clusters representing cells from the ureter and ionocytes (Fig. 1C, S1C) as indicated by expression of gene markers for ionocytes/intercalating cells in one group; these were absent in the other group (Fig. 1D, S1D, and S3). These clusters were manually separated into cluster 6a and 6b, with 6b expressing the ionocyte markers. These cell likely remained one cluster as there is only a small number of cells present in each subcluster (6a = 62 cells 6b = 17 cells).

### Comparison of Xenopus pronephric and mouse metanephric segmentation

In order to identify the cell types within the *Xenopus* pronephros, we utilized multiple approaches to compare the *Xenopus* cells to the mouse metanephric kidney data set produced by Ransick et al 2019. These data include male and female mouse samples derived from distinct anatomical regions along the corticomedullary axis of the metanephric kidney and identified 32 cell types within the nephric lineage, numbered 1-32 ^34^ (Fig. 3C, 5B-D).

**Figure 3:**
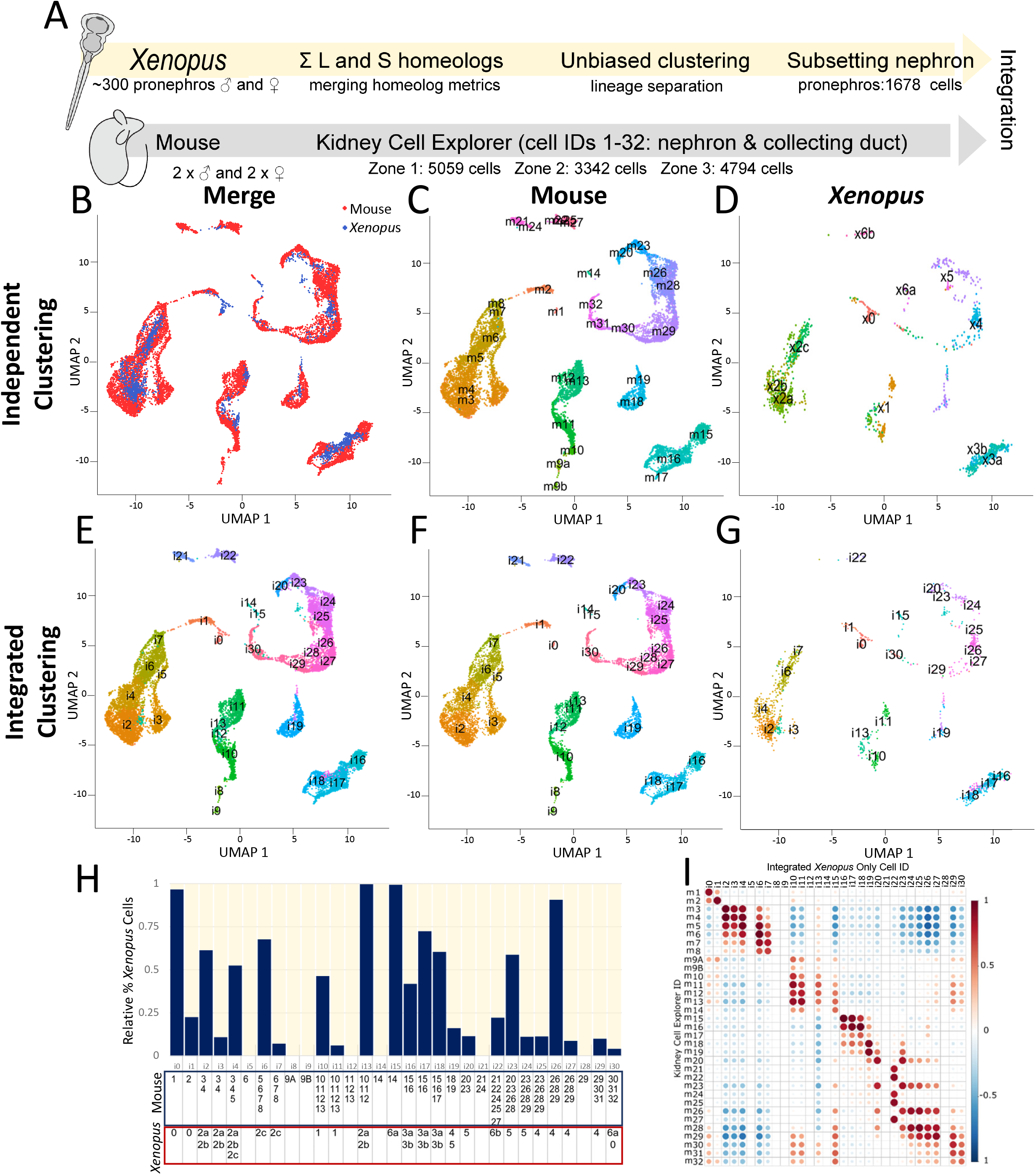
Comparison of cell types from the *Xenopus* and adult mouse nephron. **A.)** Description of methodology to generate integrated data sets. **B.)** UMAP generated from *Xenopus* pronephric and mouse metanephric cells. **C,D.**) UMAP with clusters identified by independent clustering. Mouse clusters start with an “m” *Xenopus* clusters start with an “x”. **E-G.**) Unbiased clustering based upon the integrated dataset. Integrated clusters start with an “i”. **H.)** Percentage of cells for each cluster that derive from each of the Mouse and *Xenopus* samples. Mouse colors refer to which kidney layers the cells derive from. Blue is cortex, yellow is outer medulla, and red is inner medulla. **I.)** Pearson’s correlation between the previously defined mouse Kidney Cell Explorer clusters <https://cello.shinyapps.io/kidneycellexplorer/> and the *Xenopus* cells as clustered within the integrated data set.

We performed multiple assays to compare *Xenopus* pronephric to the mouse metanephric cells. First, we utilized Seurat to integrate the two data sets and independently clustered this complied set (Figure 3), creating a new integrated cluster identity labeled with an “i” that is distinct from the original cell clusters identities in the mouse “m” or *Xenopus* “x”. Integrated clusters that were primarily composed of mouse or *Xenopus* cells are not significantly similar, while clusters that contain a significant number of both *Xenopus* and mouse cells indicate they are a similar cell type (Fig. 3H). To evaluate how well these cell types associate with each other, a Pearson correlation coefficient was calculated between the mouse Kidney Cell Explorer cell types and the integrated clusters (Figure 3I). However, a relative Pearson correlation coefficient can create false associations if the *Xenopus* cells do not align well with any of the mouse cells. To circumvent this problem, the average expression level of all the genes enriched in the *Xenopus* clusters within each mouse nephron segment was calculated (Fig. 5A). Given that seven cell clusters were identified in our primary analyses of cells in the *Xenopus* pronephros, a lower number than that identified in the mouse, cell identities from the mouse kidney cell atlas were grouped as follows: (Fig. S3C, 5D): Podocytes (1) Parietal Epithelium (2) Proximal Tubule (3-8) Loop of Henle (9a-14) Distal Tubule (15-19) Principal cells (20, 23, 26, 28, 29) Intercalated cells (21, 22, 24, 25) Duct cells (30-32). For all the genes enriched in each *Xenopus* segment, the average expression of these genes in mouse was calculated. Finally, a functional correlation between the nephron segments between the species was generated by specifically comparing genes that perform specific functions along the nephron (Fig. 4)

**Figure 4:**
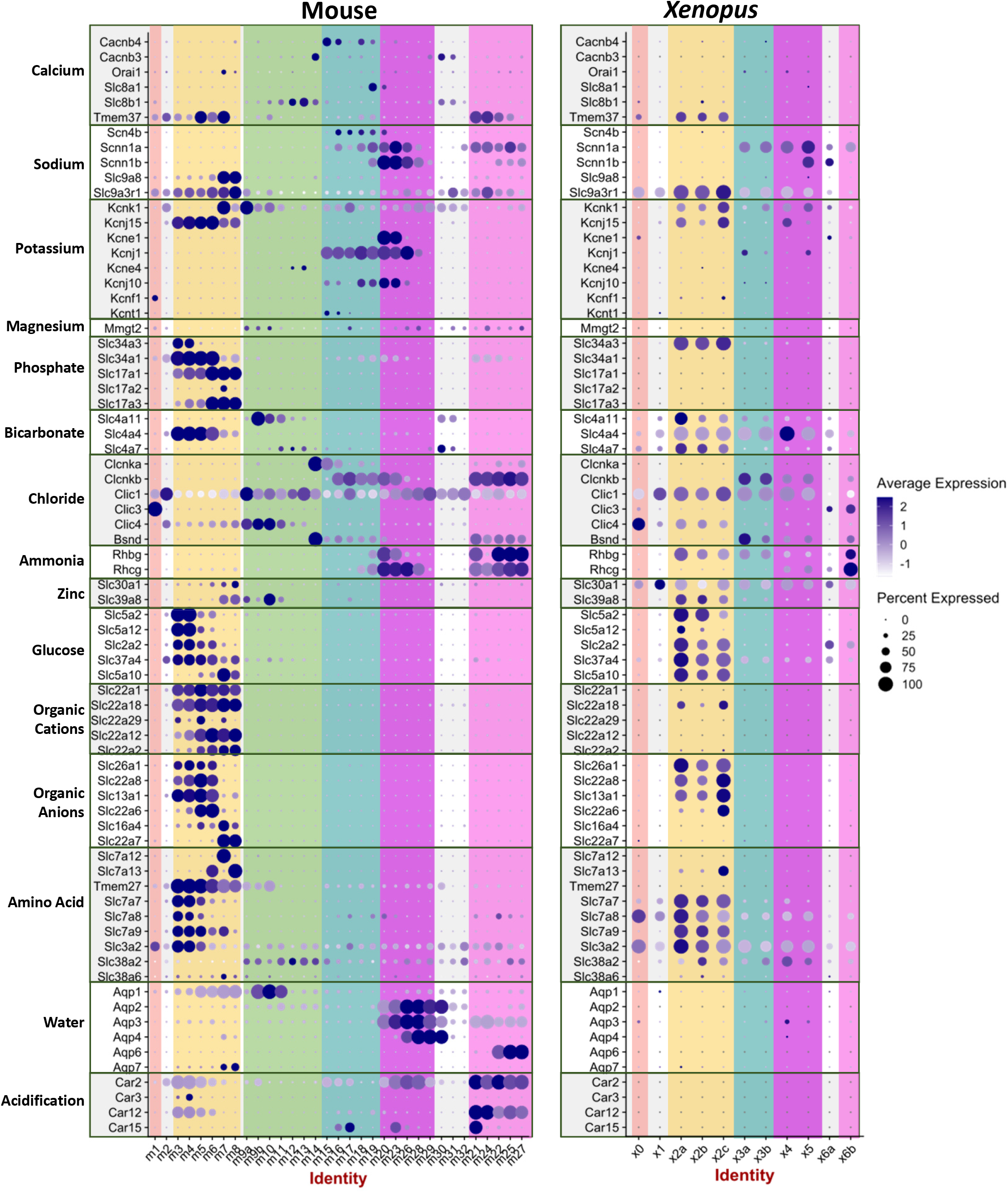
Expression patterning of genes involved in renal function are largely conserved. Dotplot of genes that are found in both mouse and *Xenopus* and play a role in kidney function.

**Figure 5:**
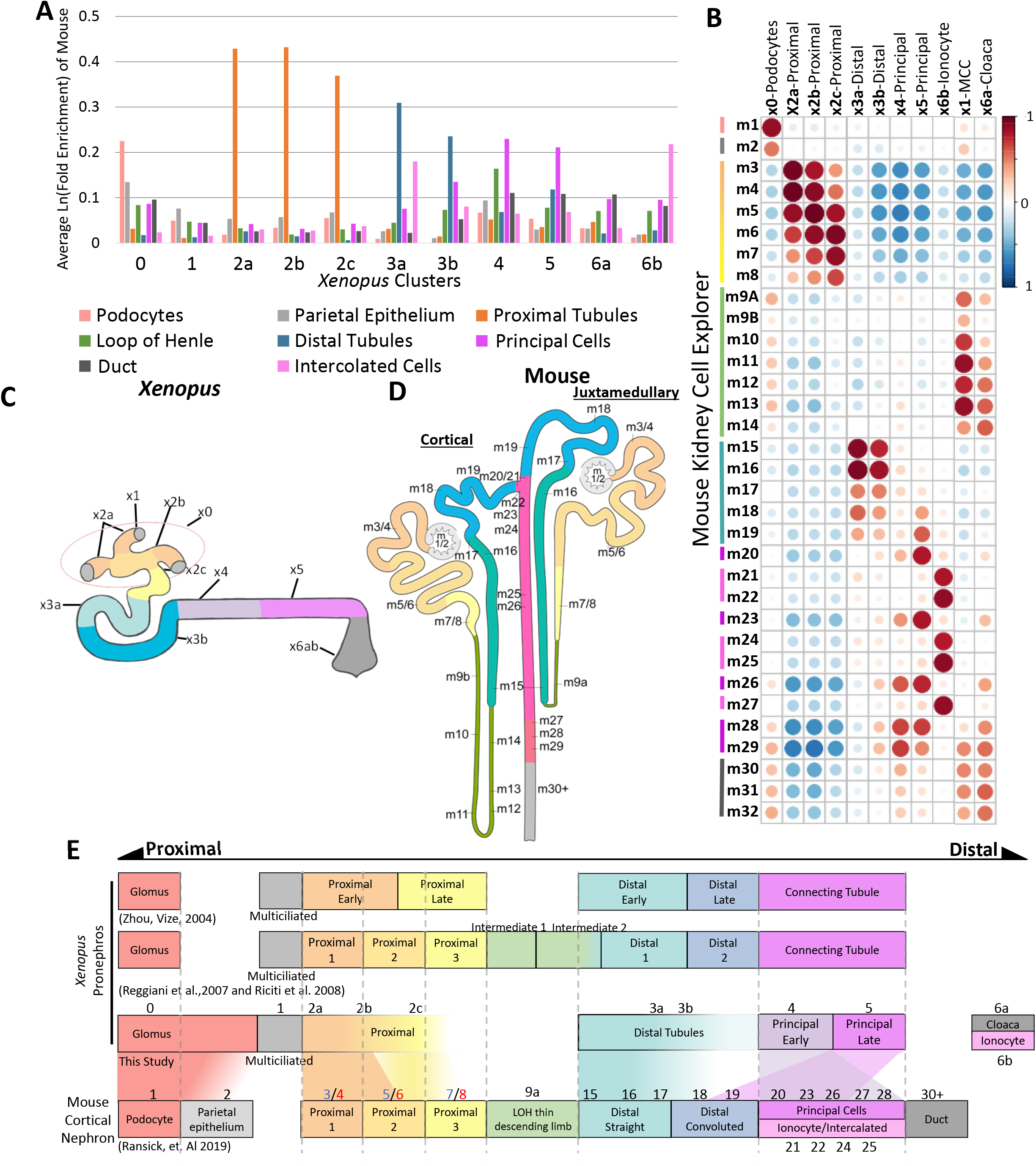
Model of how the *Xenopus* nephron segments maps onto the defined mouse nephron segments. A.) Average expression of mouse genes found enriched in each of the *Xenopus* segments. Lists of genes significantly enriched for each *Xenopus* cluster were identified. Using mouse single-cell data published by Ransick, et. al 2019, the relative expression for each gene was identified for each of the mouse nephron segments. Average expression levels of these genes are indicated. **B.)** Pearson’s correlation between *Xenopus* and Mouse clusters. **C-D.)** Diagrams showing how the *Xenopus* kidney aligns with the previous models of the *Xenopus* pronephros ^21,22^ as well as the mouse cortical nephron.

Using these methods, a number of the pronephric segments mapped cleanly onto the mouse kidney, such as the glomus/podocytes (0), proximal tubules (2a-c), Distal tubules (3a,b) and ionocytes/intercalating cells (6b) (Fig. 3C). Both groups 4 and 5 associated the best with the cells of distal convoluted tubules and the principal cells of the duct. The *Xenopus* principal cells also share similar expression patterns as those in the mouse. The more proximal cells (2a-b) best associate with the early proximal cells of the mouse (3/4), while the later proximal cells (2c) in *Xenopus* best correlate with the middle proximal segment in mouse.

Other segments did not map as cleanly onto the mouse kidney, such as the multiciliated cells (1) and cloacal cells (6a) There is some weak correlation with the cloaca and collecting duct cells when compared to the mouse, suggesting it may be some of the most distal cells of the kidney. This cluster expresses markers indicating similarity to the most distal duct and the bladder such as the uroplakin (*Upk*) family of genes (Fig. 1D S1C, S4).

Through our single-cell transcriptomic analysis, we have refined the model of the *Xenopus* pronephros as compared to published models based upon *in situ* analysis (Fig. 5D) (Raciti et al., 2008; Reggiani et al., 2007; Zhou and Vize, 2004). forming a model for the frog nephron as compared to the mouse cortical nephron (Fig. 5C).

### Single-cell profiling and clustering of kidney associated cells

Dissection of the pronephros resulted in isolation of several cell types other than nephric epithelial cells. We have separated them into three groups: neural crest, vasculature, and mesenchyme. It is unclear as to whether these cells surrounding the nephron play a role in supporting its function. However, gene pattering does not appear similar to these cell types associated with the kidneys in mice^39,40^.

Multiciliated cells associated heavily with mesenchymal cells but are physiologically part of the nephrostome of the kidney (Fig. 6D, S2C). Therefore, this group was included in both the mesenchymal group as well as the kidney group. Each group was subclustered and independently processed.

**Figure 6:**
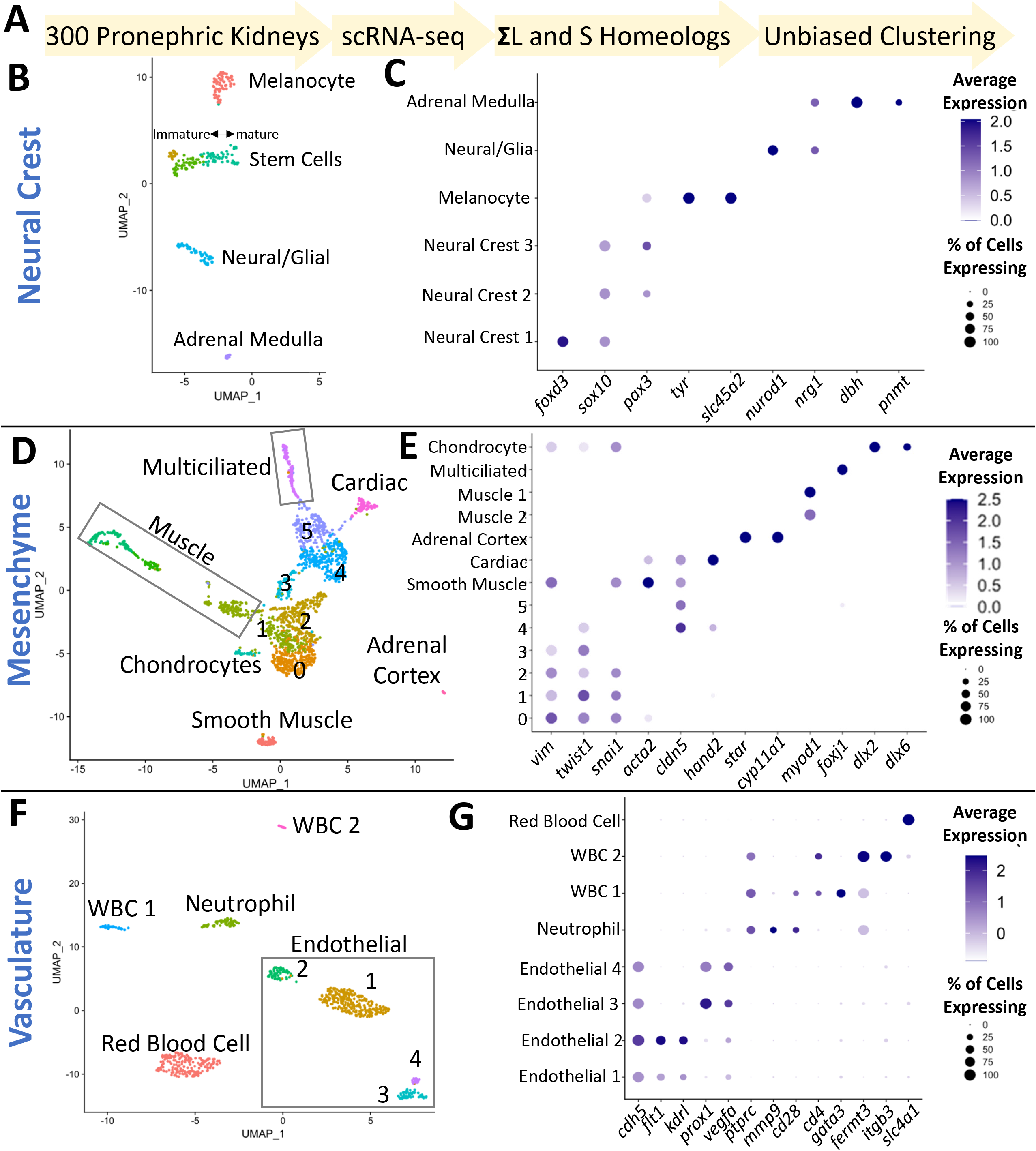
Identity of other cell types isolated with the *X. laevis* kidney. UMAP showing subclustered cell types: neural crest **(A)**, mesenchyme **(C)**, and vasculature **(E)**. Dotplot showing the expression of validated markers for each cell type from neural crest, mesenchymal, and vasculature subsets **(B,D,F).**

The neural crest population falls into four major groups: neurons (*neurod1* and *nrg1*) ^41,42^, melanocytes (*tyr* and *slc45a2*) ^43,44^, adrenal medulla (*dbh* and *pnmt*) ^45^, and undifferentiated cells (*foxd3, sox10, pax3*) ^46,47^ (Fig. 6B, S2A). The undifferentiated neural crest cell cluster separated into three groups by unbiased clustering, with the group labeled Neural Crest 1 being the more immature cells as they express the transcription factor *foxd3*, which functions to inhibit differentiation by repressing factors such as *pax3* ^48–50^. As expected, the more differentiated cell types such as neurons have lost expression of these canonical early neural crest markers apart from the melanocytes, which retained *pax3* expression ^46^.

The largest number of cells isolated fall into the mesenchymal category. We have defined these cells by UMAP analysis. These cells associate with a central group of cells that express the mesenchymal markers *vim, snai1*, and *twist1* (Fig. 6D,E, and S2C,D) ^51,52^. However, a cell classification could not be identified for a number of these groups. This is likely because they are undifferentiated cells and do not strongly express markers of mature cell types. Therefore, these cell clusters were numbered mesenchymal 1-5. There were a number of cells types which could be identified using well established markers, such as: smooth muscle (*acta2/sma*), cardiac (*cldn5* and *hand2*), adrenal cortex (*star* and *cyp11a1*), skeletal muscle (*myod1*), and chondrocytes (*dlx2* and *dlx6*). Though the multiciliated cells associated with the mesenchymal group by UMAP, they are also a part of the nephrostome of the kidney. Therefore, they are included in both groups for analysis.

Cells clustered in the vascular group include both the vasculature and blood cells. Characterized markers were used to distinguish endothelial cells (*cdh5/ve-cadherin*), white blood cells (ptprc/*cd45*) ^53^, and red blood cells (*Slc4a1*) ^54^ (Fig. 6F,G S2E,F). The vasculature separated into four groups labeled endothelial 1-4. Endothelial clusters 1 and 2 share markers with blood vasculature such as *flt1/vegfr1, and kdrl/vegfr2* ^55^, while clusters 3 and 4 share markers with lymphatic vessels such as *prox1* and *vegfa* ^56^.

Three categories of Ptprc/CD45+ white blood cells (WBC) were identified, neutrophils and two other white blood cell types (Fig. 6F,G S2E,F). The identity of two of the white blood cell groups are largely unknown as many of the markers used for identification in mouse and humans are not conserved in *Xenopus*. By GO ontology analysis < http://geneontology.org/> comparing the gene list to human, WBC group 1 does share some characteristics of lymphocytes such as *gata3* and *il4*, and group 2 share some gene expression with platelets *itgb3* and *fermt3* ^57^. Though they were significantly enriched for the indicated cell identity, the number of genes shared was still low 4 and 6 respectively.

The adrenal glands develop from two separate embryological tissues: the medulla is derived from neural crest cells originating in proximity to the dorsal aorta (Fig. 6B, S2B), while the cortex develops from the intermediate mesoderm near the proximal tubules of the kidney (Fig 6F, S2F) ^58,59^. In *Xenopus*, no research has been done looking at the development of the adrenal gland prior to metamorphosis; therefore, these cell types were identified using the human expression profile <https://www.proteinatlas.org/humanproteome/tissue/adrenal+gland> ^60^. The expression data in this study provides markers that can be used to track and visualize the development of the adrenal gland in *Xenopus*. We were able to identify the adrenal medulla as part of the neural crest type and the adrenal cortex as part of the mesenchymal cell types.

## Discussion

The goal of this study was to identify all the cell types in *X. laevis* pronephric kidney and to understand the patterning of the nephron in relation to the mammalian kidney. Seven cell types were characterized in the *X. laevis* kidney: glomus, multiciliated cells, proximal, distal early, distal late, principle like, cloaca and ionocytes.

### Technical choices of stage and genome

Different stages are commonly used to study kidney development in *Xenopus*. Stages 12.5-20 provide a window to evaluate the initial inductive events that establish the kidney progenitors ^43,61,62^. At stage 30, cellular movements and epithelization leads to the formation the epithelial tube ^22,63^. By stage 37, the nephron begins to function, with specialized segments that reabsorb nutrients as fluid starts to flow through the kidney ^22,64^. The purpose of this study is to identify cell types in the functional kidney to build upon existing models of nephron segmentation in *Xenopus* and to evaluate the conservation of this patterning in the mammalian metanephric kidney. To evaluate the cell types within the functional nephron, stage 40 kidneys were isolated for single-cell sequencing, as we predict at this stage all the cell types have been established. Additionally, there are technological advantages to choosing stage 40. Therefore, stage 40 was selected as the optimal time point to address this question.

The *X. laevis* genome is still being annotated with provisional or unannotated genes. Currently, the official genome published in 2017 is continuing to be annotated ^38^. In recent years, increased efforts have been set forth to bring both the *X. laevis* and *X. tropicalis* genome annotations up to the quality of other commonly used model organisms ^65^. Periodically, an unofficial pre-release genome is made available to the community with greatly improved annotation. Therefore, for the purposes of this study, the pre-release genome that was uploaded in January of 2020 was used. The genes that are still under review are labeled with a “Loc” number and are labeled as provisional genes.

### Comparison of different Xenopus models of the kidney

Our single-cell analysis of the *Xenopus* nephron builds upon valuable positional *in situ* evaluation of nephric gene expression to generate models of nephron segmentation. Our results largely substantiate both the models proposed by Zhou and Vize 2004, and Raciti, et al., 2008, with some exceptions (Fig. 5D). The Zhou and Vize model identified two proximal segments, while the Raciti et al. model suggested three segments. Our new single-cell data indicate that here are a few genes, such as *slc6a19* and *slc5a8*, which create an intermediate proximal region. By *in situ*, there appears to be a clean division between the three segments of the proximal tubule; however, single-cell analysis indicates a gradient from of one cell type to another. This may result from a number of factors: First, *in situ* analysis is not extremely quantitative, and often only the strongest expressing cells can be detected, hiding sharp gradients and weaker expressing cells. Conversely, single-cell analysis is more quantitative, picking out more intermediately expressing cells. Secondly, *in situ* evaluates the expression a single gene in a single embryo, while our analysis looks at many genes from many embryos. Therefore, if the gene expression pattern of a particular marker transitions at a slightly different position or varies slightly different from embryo to embryo, the UMAP will mix cell populations around the transition resulting in cell type separation that is more gradual by single-cell analysis.

The Raciti et al. model suggests that the cell type 3a-b is part of the loop of Henle (LOH). The majority of genes analyzed to support this conclusion, including *cldn8*, are in both the Loop of Henle and distal tubules of mice. Another gene assessed that is unique to the mouse loop of Henle is *clcnka*. However, *Xenopus* only possesses a single *clcnk* gene annotated *clcnkb*. In both mouse and *Xenopus clcnkb* is expressed in the more distal tubules such as the connecting tubules and collecting duct. Thus, the data do not confirm or refute the existence of an intermediate tubule in *Xenopus*. Additional evidence for the Loop of Henle in *Xenopus* is the expression of the three Iroquois genes (*irx1, irx2*, and *irx3*) in *Xenopus* at the distal end of the proximal tubules and the proximal end of the distal tubules (clusters 2c and 3a). By *in situ* analysis, these genes look like they are in the LOH in mouse; however, single-cell analysis indicates that these gene are not only expressed in the LOH, but are also expressed elsewhere in the kidney. Although these data may indicate the presence of LOH cells in the frog pronephros, other explanations are possible. One of the main functions of the LOH is to reabsorb water. This is done with the aid of aquaporins. *aqp11 and ap3 are* the only aquaporin expressed in the *Xenopus* kidney, and it is only weakly expressed in the proximal tubules and early principal respectively. By Pearson’s correlation the multiciliated cells (1) and cloacal cells (6b) best correlate with the LOH and duct cells (Fig. 5B). However, this is likely an artifact of not having a similar cell type present in mouse. Looking at average gene expression (Fig. 5A) there is no enrichment of gene expression. Unlike the pronephric and mesonephric kidney the metanephric kidney does not have multiciliated cells. Additionally comparing expression of enriched genes across species does not show an enriched expression of LOH genes in these clusters. Therefore, we do not believe that there is an equivalent to the LOH in *Xenopus*. We also would not predict a need for a LOH to concentrate urine, as *Xenopus* are freshwater animals and water is not a limiting nutrient.

Mammalian metanephric kidneys contain two types of nephrons cortical and juxtamedullary ^66^. The cortical nephrons have a shorter LOH that only extends into the outer zone of the renal medulla, and primarily functions to reabsorb nutrients. Because we do not find strong evidence for the presence of a LOH in *Xenopus*, the pronephric kidney most closely resembles the cortical nephrons of the metanephric kidney.

Both published models indicate the existence of a connecting tubule; however, the mouse connecting tubule contains two cell types the principal cells and two types ionocytes/intercalating cells. We find the cells in the connecting tubule region (Cluster 4 and 5) of the *Xenopus* kidney most closely resembles the principal cells of the mouse. Ionocytes are cells that regulate pH. In the mouse kidney, there are two types of intercalating cells type A pumps protons largely through V-type atpases, while type B pumps bicarbonate largely through the Slc26a4 family of transporters. We are able to detect one type of intercalating cells that are mostly associated with the cells of the cloaca (6b). This cell type expresses both types of transporters suggesting it functions as both Type A and type B ionocytes however this might be an artifact of the few cells we were able to sequence.

By UMAP analysis, we observe that the multiciliated cells show a spectrum of cells that are more or less associated with the undifferentiated mesenchymal cells. This may indicate that the kidney is still adding multiciliated cells or there is a large turnover of this cell type. Around the stage in which we isolated kidneys for this study the multiciliated cells of the epidermis are undergoing removal ^67^. This occurs by two processes, apoptosis and cellular reprogramming to a different cell fate. As we do not see a strong expression of p53 regulated genes in any tissue isolated in this study, we don’t predict that this is due to turnover of ciliated cells. At this point in development, the largest growth in tubules appear to be in the proximal tubules; therefore, it is possible that the kidney is still adding multiciliated cells ^64^.

Three families of genes are missing from the *Xenopus* kidney (Fig. 4). These include genes involved in transport of water, transport of organic cations, and carbonic anhydrases which are involved in pH balance. In the early tadpole much if the kidney function is likely carried out through the epidermis prior to the formation of the functioning nephron. Therefore, much of the pH regulation in the early *Xenopus* tadpole is performed in the epidermis which is covered with ionocytes which can pump protons out of the organism this is what likely accounts for loss of these gene families within the kidney. *Xenopus* is also a freshwater organism, so removal of water is more essential than reabsorption as they are living in a hypotonic environment with water constantly entering.

Surprisingly our podocyte population shares gene expression with that of the parietal epithelium (PE) (Fig 5A,B). In Mice these cells develop from the same mesenchymal pool^31,68^ and it has been speculated that the PE functions as the stem cell niche for podocyte regeneration^69^. As the Xenopus pronephros does not contain a clear PE cell type it is possible that the podocytes perform both PE and podocyte function.

### Conclusion

In this study we have identified the cell types of the kidney, generated a cell expression profile for each of these types, and successfully mapped the *X. laevis* nephron segments onto the mouse kidney. We have also isolated cells that physically associated with the kidney cells and have generated an expression profile for each of these cell types. These data can be used to find markers that will aid in future research to facilitate the development of disease models.

## Materials and methods

### Xenopus laevis care

Wild type *X. laevis* adult males and females were purchased from Nasco (LM00713M and LM00531MX). All embryos were grown in 1/3X MMR (33 mM NaCl, 0.66mM KCl, 0.33 mM MgSO_4_, 0.66 mM CaCl_2_, 1.66 mM HEPES pH7.4). This protocol was approved by the UTHealth’s Center for Laboratory Animal Medicine Animal Welfare Committee, which serves as the Institutional Animal Care and Use Committee (protocol #: AWC-19-0081).

### Kidney Isolation

Wild type *X. laevis* were grown to stage 40 using standard protocols ^70^. Animals were anesthetized 0.02% Benzocaine (Spectrum BE130) diluted in 1xMMR (100mM NaCl, 2mM KCl, 1mMMgSO_4_, 2mM CaCl_2_, 5mM HEPES pH7.4). Manipulations were carried out on 2% agar melted in 1xMMR. Dissections were performed as demonstrated in movie S1. Kidneys were moved using a glass pipette and stored in a 1.5ml tube 1xMMR until disassociation. Approximately 300 kidneys were dissected for further processing.

### Dissociation of kidney cells

Kidneys were washed twice in 1xMMR and allowed to settle by gravity. Liquid was removed and replaced with 50:50(1xTrypLE (Gibco 12605010):1xMMR) supplemented with 1mM L-cystine (Alpha Aesar A10435) + 10U/ml Papain. The 1.5ml tube was gently inverted once every 10 minutes and allowed to incubate on its side at room temperature until kidneys started to disassociate (~1.5hrs). The tube was topped off with 45:45:10 1xMMR:L15:FCS to inactivate the papain and strained threw a 70um sieve (VWR 10199-656). The sieve was washed with 25ml 45:45:10 1xMMR:L15:FCS. Cells were pelleted 500xg for 5min. Supernatant was removed and ~500ul 1xMMR was added to the tube and cells were gently resuspended by shaking, then moved to a 1.5ml tube containing 1:1 1xMMR:Optiprep (Cosmo Bio Usa Inc AXS1114542) with a glass pipette. Cells were centrifuged 1000xg for 5min and the upper and interphase layer was moved to a new tube with a glass pipette, leaving yolk pellets in the denser lower layer. Cells were washed 2x with PBS by centrifugation 500xg 5min to remove residual Optiprep and to prevent clumping.

### Quality Control and Single-Cell Sequencing

scRNA seq was carried out at the ATGC genomics core at MD Anderson Cancer Center. Cells were stored on ice in PBS during quality control. Cells labeled with Trypan blue and visualized on a hemocytometer. Multiciliated cells were motile and there were minimal signs of apoptosis. Additionally, there was approximately one yolk pellet per cells. 10,000 cells were used for 3’ scRNAseq using the 10x genomics platform. cDNA library was run on Illumina NextSeq500 system.

### Data Processing and Statistics

Sequences were processed using the CellRanger 3.1.0 platform. A compatible genome was created using the pre-release (01-31-2020) *Xenopus laevis* 9.2 genome available at the Xenbase FTP server http://ftp.xenbase.org/pub/Genomics/JGI/Xenla9.2/pre-release/. Seurat V3.2.3 was used to perform cluster analysis. As the *X. laevis* genome does not have the mitochondrial RNAs labeled in the reference genome cells with greater than 12% mitochondrial 12S and 16S ribosomal RNA’s were discarded as poorly sequenced cells. As an additional cutoff, cells with greater than 90,000 reads/cell or lower than 3000 reads/cell were removed. Normalization and clustering were performed as described https://satijalab.org/seurat/^71^.

For mouse single-cell analysis data was downloaded from NCBI BioProject Accession PRJNA532850 ^34^. All 12 samples were processed and combined through Cell Ranger 3.1.0 using the mouse mm10 reference genome, and clustering was done in Seurat 3.2.3.

Unless otherwise stated, an enriched gene is defined as a gene that has an average log fold change (avg_logFC) of 0.5 and adjusted p value(p_val_adj) < 1E-5 will within the cluster. Because eSeurat uses natural log for its fold change calculations, a value of 0.5 is approximately equivalent to a 1.6-fold enrichment. A less stringent cutoff was used to generate the supplemental datasets, however only genes that fit the criteria were used for analysis.

Relative cell numbers were calculated as (# *Xenopus* cells in cluster / total # *Xenopus* cells) / (# mouse cells in cluster / total # mouse cells)

## Acknowledgements

We appreciate the helpful suggestions and advice throughout this project from the members of the laboratories of R. K. Miller, P. D. McCrea, J. Park, as well as M. Kloc. A special thanks to Dr. Yoshihiro Komatsu for the use of equipment in this project. We thank the animal care technicians and veterinarians, including J.C. Whitney and T.H. Gomez who took care of the animals. The scRNA sequencing was performed using the 10X genomic platform ran and maintained by the advanced technology genomics core (ATGC) at MD Anderson Cancer Center, and we are grateful to David Pollock for his technical support.

## Author contributions

This project was initially conceptualized by MEC and RKM. Kidney dissections performed by MEC, BDD, and VKS. Conditions for disassociation were tested and troubleshooted by MEC and BDD. Data analysis was performed by MEC, MAA and NOL with the help of MPC and JC. Graphical design of Fig. 5 generated by BLW. Writing and figures were initially assembled by MEC. Project administration as well as data and manuscript evaluation were carried out by MEC and RKM.

---

**Figure S1:**
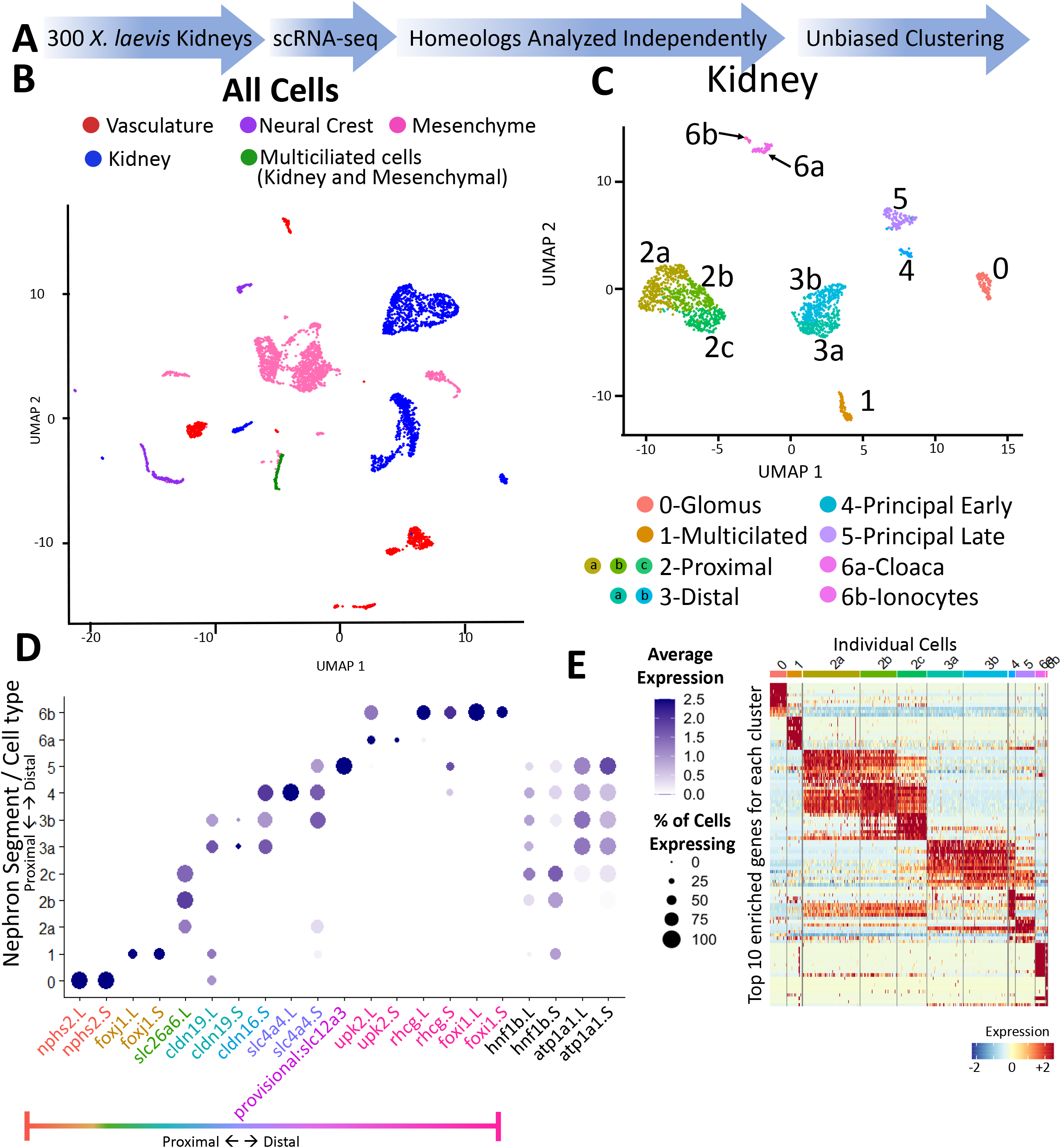
Clustering of cells after combining homeologs. **A.)** Description of methodology to generate integrated data sets. **B.)** UMAP of all cells isolated. **C.)** UMAP of subclustered kidney cells. **D.)** Dotplot showing expression of validated mouse markers in the *X. laevis* pronephros **E.)** Heatmap showing the expression of the top 10 most enriched genes for each cluster. Rows indicate gene expression and columns indicate a single cell.

**Figure S2:**
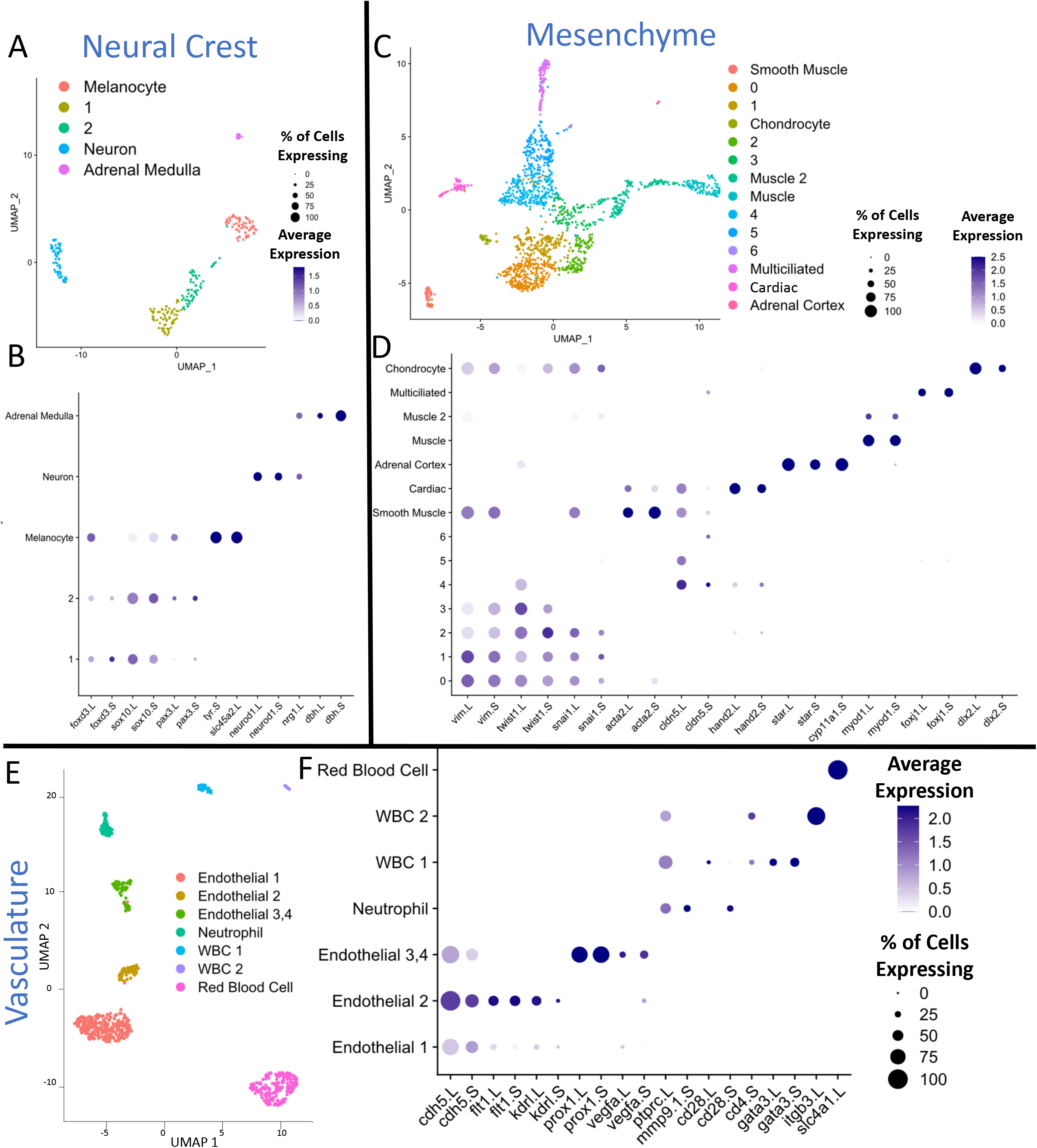
Clustering of non-nephron cells prior to the combining of homeologs. Cells were subclustered from the total cells (Fig. S1) into neural crest, mesenchyme and vasculature. **A, C, E.)** UMAP of indicated subpopulation of cells. **B, D, F.)** Dotplot showing represented genes for each cell type.

**Figure S3:**
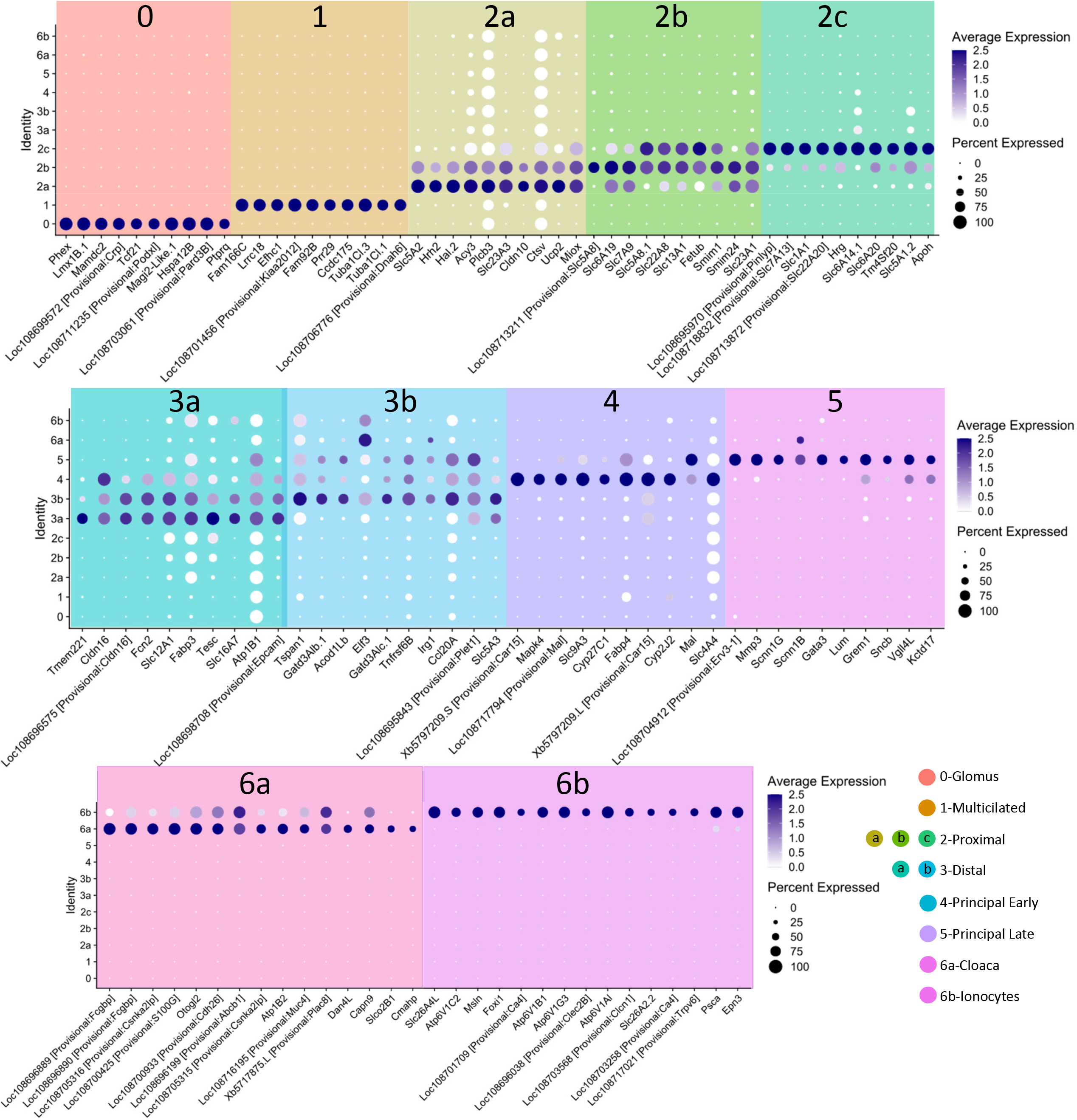
Dotplot showing expression of the top 10 most enriched genes for each cluster.

**Figure S4:**
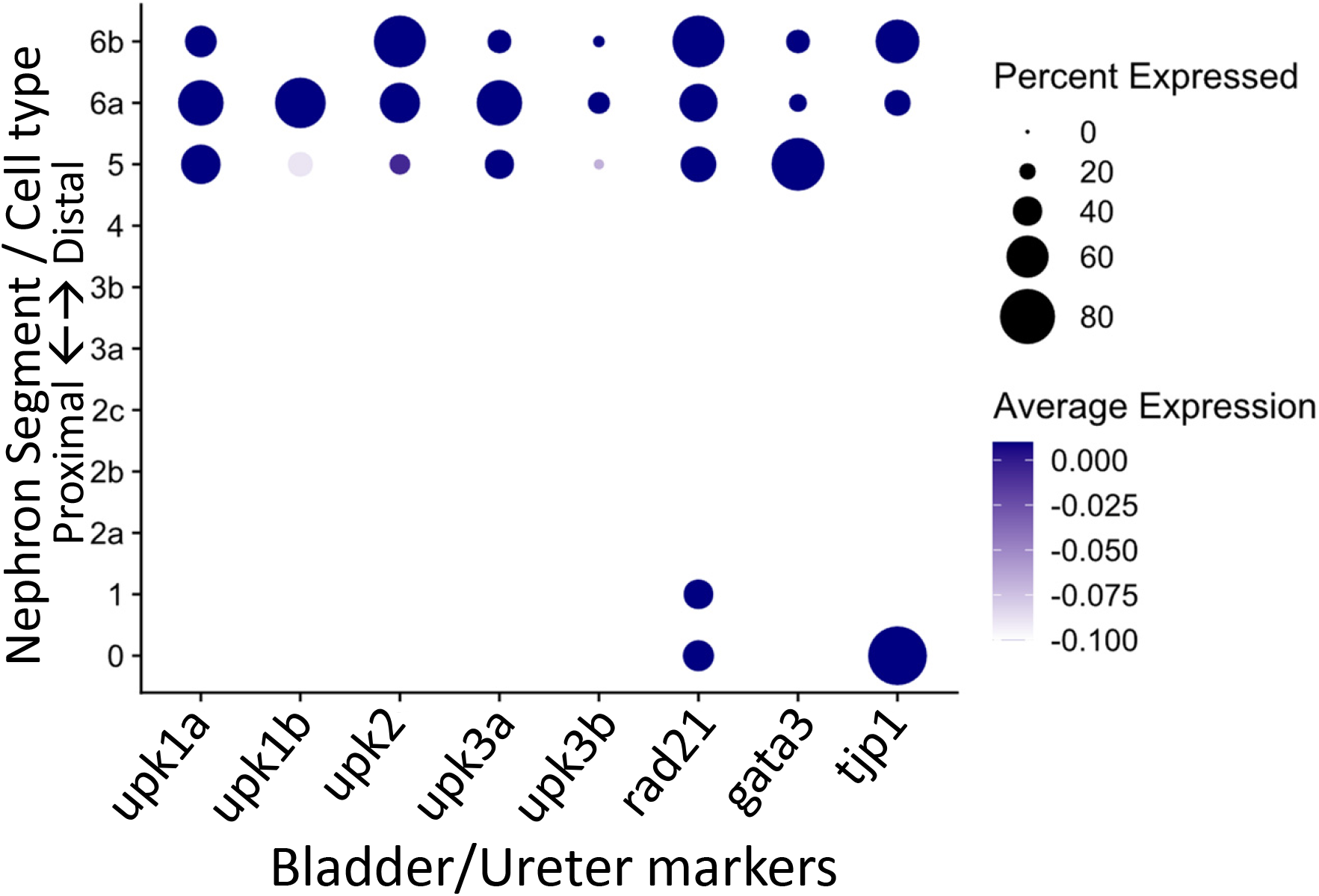
Dotplot showing expression of published genes for the ureter and bladder. Expression of each cell for indicated gene as displayed by UMAP. Darker blue dots indicate higher expression.

**Movie S1:** Movie showing the dissection of the *Xenopus* kidney played at 2x speed.

**Table S1:** List of genes enriched prior to grouping homeologs.

**Table S2:** Table of gene counts and cell labels after grouping homeologs for each cell.

**Table S3:** List of genes enriched after to grouping homeologs.

